# 4CAC: 4-class classifier of metagenome contigs using machine learning and assembly graphs

**DOI:** 10.1101/2023.01.20.524935

**Authors:** Lianrong Pu, Ron Shamir

**Affiliations:** Blavatnik School of Computer Science, Tel Aviv University

## Abstract

Microbial communities usually harbor a mix of bacteria, archaea, plasmids, viruses, and microeukaryotes. Within these communities, viruses, plasmids, and microeukaryotes coexist in relatively low abundance, yet they engage in intricate interactions with bacteria. Moreover, viruses and plasmids, as mobile genetic elements, play important roles in horizontal gene transfer and the development of antibiotic resistance within microbial populations. However, due to the difficulty of identifying viruses, plasmids, and microeukaryotes in microbial communities, our understanding of these minor classes lags behind that of bacteria and archaea. Recently, several classifiers have been developed to separate one or two minor classes from bacteria and archaea in metagenome assemblies, but none can classify all of the four classes simultaneously. Moreover, existing classifiers have low precision on minor classes. Here, we developed a classifier called 4CAC that is able to identify viruses, plasmids, microeukaryotes, and prokaryotes simultaneously from metagenome assemblies. 4CAC generates an initial four-way classification using several sequence length-adjusted XGBoost models and further improves the classification using the assembly graph. Evaluation on simulated and real metagenome datasets demonstrates that 4CAC substantially outperforms existing classifiers and combinations thereof on short reads. On long reads, it also shows an advantage unless the abundance of the minor classes is very low. 4CAC runs 1-2 orders of magnitude faster than the other classifiers. The 4CAC software is available at https://github.com/Shamir-Lab/4CAC.

## 1 Introduction

Microbial communities in natural and host-associated environments are often dominated by bacteria and coinhabited by archaea, fungi, protozoa, plasmids, and viruses [25]. Changes in microbiome diversity, function, and density have been linked to a variety of disorders in many organisms [26, 10]. As the dominant group of species in microbial communities, bacteria have been widely studied. Taxonomic classification tools [43, 42] and metagenome binning tools [24, 23, 14, 44] were proposed to detect bacterial species present in a microbial community directly from reads or after assembling reads into contigs. It is known that the specific composition and abundance of certain bacterial species affect their host’s health and fitness [6, 20, 27]. In contrast, our understanding of plasmids, viruses, and microbial eukaryotes largely lags behind, due to their lower abundance and the difficulty of detecting them in microbial communities [4, 21]. Recent studies revealed that viruses and plasmids play important roles in horizontal gene transfer events and antibiotic resistance [7, 40, 22, 39], and microbial eukaryotes have complex interaction with their hosts in both plant- and animal-associated microbiomes [21, 29]. To better understand the species composition and the function of each species in microbial communities, classifiers that can identify not only bacteria but also the other members of a microbial community are needed.

Many binary and three-class classifiers have been developed in recent years for separating viruses and plasmids from prokaryotes (bacteria and archaea) in microbial communities. VirSorter [36], DeepVirFinder [35], VIBRANT [16], and many other classifiers [12, 3] were designed to separate viruses from prokaryotes. Plasmid classifiers, such as PlasFlow [18], PlasClass [30], Deeplasmid [1], and Platon [37] were developed to separate plasmids from prokaryotes. As both viruses and plasmids are commonly found in microbial communities, three-class classifiers, such as PPR-Meta [9], viralVerify [2], 3CAC [33], and geNomad [8] were proposed to simultaneously identify viruses, plasmids, and prokaryotes from metagenome assemblies. In contrast, microbial eukaryotes, such as fungi and protozoa, are integral components of natural microbial communities but were commonly ignored or misclassified as prokaryotes in most metagenome analyses. More recently, EukRep [41], Tiara [15], and Whokaryote [32] were proposed to distinguish microeukaryotes from prokaryotes. However, even though prokaryotes, microeukaryotes, viruses, and plasmids are present in most microbial communities, to the best of our knowledge, there are still no four-class classifiers that can simultaneously identify and distinguish all of them. (A recent preprint reported a five-way classifier, DeepMicrobeFinder [13], but the code provided on GitHub was not functional.) Moreover, most classifiers ignore the fact that microbial communities are dominated by bacteria, and have low precision on the minor classes, such as viruses, plasmids, and microeukaryotes [35, 33].

In this work, we present 4CAC (4-Class Adjacency-based Classifier), a fast algorithm to identify viruses, plasmids, microeukaryotes, and prokaryotes simultaneously from metagenome assemblies. 4CAC generates an initial classification using a set of XGBoost models trained on known reference genomes. The XGBoost classifier outputs four scores for each contig to indicate its confidence of being classified as a virus, plasmid, prokaryote, or microeukaryote. To assure high precision in the classification of minor classes, we set higher score thresholds for classifying minor classes compared to prokaryotes. Subsequently, inspired by 3CAC, 4CAC utilizes the adjacency information in the assembly graph to improve the classification of short contigs and of contigs with lower confidence by the initial classification. Evaluation of 4CAC against combinations of existing classifiers on simulated and real metagenome datasets demonstrates that 4CAC has substantially better performance on short reads. On long reads, it also shows an advantage unless the abundance of the minor classes is very low.

## 2 Results

### 2.1 The 4CAC algorithm

To understand the species present in a microbial community, the common practice is to first assemble the sequencing reads into longer sequences called *contigs*, and then classify these contigs into classes. Broadly used metagenome assemblers, such as metaSPAdes [28] and metaFlye [17], use assembly graphs to combine overlapped reads (or k-mers) into contigs. Nodes in an assembly graph represent contigs and edges represent sequence overlaps between the corresponding contigs. Most of the existing classifiers take contigs as input and classify each of them independently based on their sequence. Our recent work on three-class classification demonstrated that neighboring contigs in an assembly graph are more likely to come from the same class and thus the adjacency information in the graph can assist the classification [33]. Therefore, here we introduce 4CAC, a four-class classifier that combines machine learning methods with assembly graph neighborhood information to classify each contig as virus, plasmid, prokaryote, microeukaryote, or uncertain.

Inspired by previous studies [30, 9, 34], to assure good classification of sequences with different lengths, we constructed five XGBoost models trained on sequence fragments of length 0.5k, 1kb, 5kb, 10kb, and 50kb, respectively. The *k*-mer composition of each fragment was used as the feature vector to train the XGBoost models. Given a sequence, we calculate its *k*-mer composition and classify it with the model that matches its length. More details on the design and implementation of the XGBoost classifiers can be found under Methods.

The XGBoost classifier outputs four scores between 0 and 1 for each sequence indicating its confidence of being classified as virus, plasmid, prokaryote, or microeukaryote. Existing algorithms [9, 30, 34] usually classify a sequence into the class with the highest score by default. To improve precision, a threshold can be specified, and sequences whose highest score is lower than the threshold will be classified as “uncertain”. However, due to the overwhelming abundance of prokaryotes in the metagenome assemblies (usually ≥ 70%), a high threshold results in low recall in the classification of prokaryotes, while a low threshold results in low precision in the classification of the minor classes (virus, plasmid, and microeukaryote). Taking into consideration the class imbalance, we chose to set different thresholds for different classes. By default, a score threshold of 0.95 was set for viruses and plasmids, and no score threshold was set for prokaryotes and eukaryotes. See Section “Length-specific classification” in Methods for more explanation on the choice of the score thresholds. This results in high precision for the classification of each class while maintaining high recall for the classification of prokaryotes.

Next, we exploit the assembly graph to improve the initial classification by the following two steps. (1) Correction of classified contigs. For a classified contig *c*, if it has at least two classified neighbors and all of them belong to the same class while *c* belongs to a different class, 4CAC corrects the classification of *c* to be the same as its classified neighbors. (2) Propagation to unclassified contigs. For an unclassified contig *c*, if all of its classified neighbors belong to the same class, 4CAC assigns *c* to that class. See Section “Refining the classification using the assembly graph” in Methods for more details.

### 2.2 Simulated metagenomes

To evaluate the performance of 4CAC and existing classifiers, we simulated two short-read and two long-read metagenome datasets as follows. Prokaryotes, their co-existing plasmids, viruses, and microeukaryote genomes were selected from the NCBI GenBank Database to mimic species in a microbial community. All the genomes selected were released after December 2021, and thus they were not included in the training set of the classifier. As a *generic metagenome* scenario, we simulated reads in proportions mimicking regular metagenomic environments. As a *filtered metagenome* scenario, where reads from large genomes are filtered, the proportions were adjusted so that plasmids and viruses are enriched. The relative abundance of genomes within each class was set as in [30]. Short reads were simulated from the genome sequences using InSilicoSeq [11] and assembled by metaSPAdes. Long reads were simulated from the genome sequences using NanoSim [46] and assembled by metaFlye. Full details on the simulation and the assembly are provided in Methods. We denote by **Sim_AN** the simulation with A=S for short reads and A=L for long reads, N=G for the generic scenario and N=F for the filtered scenario. For example, Sim SF is the short read filtered scenario. Table 1 summarizes the properties of the datasets and the assemblies.

**Table 1.** Properties of the simulated and the real metagenomic datasets.

| Dataset | Read type | Number of read(M) |  |  |  | Number of assembled contigs |  |  |  | Short contigs<br>(< 1kb) |
| --- | --- | --- | --- | --- | --- | --- | --- | --- | --- | --- |
|  |  | prokaryote | eukaryote | plasmid | virus | prokaryote | eukaryote | plasmid | virus |  |
| Sim_SG | MiSeq | 56 | 24 | 10 | 10 | 15,460 | 8,112 | 1,725 | 1,275 | 5,095 |
| Sim_SF | MiSeq | 3.5 | 1.5 | 10 | 10 | 50,546 | 44,378 | 1,650 | 1,256 | 56,735 |
| Sim_LG | Nanopore | 0.56 | 0.24 | 0.1 | 0.1 | 1,575 | 148 | 193 | 202 | 125 |
| Sim_LF | Nanopore | 0.035 | 0.015 | 0.1 | 0.1 | 922 | 343 | 207 | 193 | 33 |
| Sharon | HiSeq | 106.3 in total |  |  |  | 3,097 | 533 | 87 | 21 | 1,169 |
| Tara | HiSeq | 190.7 in total |  |  |  | 16,156 | 31 | 153 | 1,270 | 11,643 |
| Oral_Nano | Nanopore | 5.6 in total |  |  |  | 9,112 | 50 | 11 | 23 | 1,888 |
| Gut_HiFi | Pacbio HiFi | 1.9 in total |  |  |  | 4,958 | 0 | 27 | 30 | 203 |

### 2.3 4CAC outperforms existing classifiers on simulated metagenomes

To evaluate 4CAC in classifying viruses, plasmids, and eukaryotes from metagenome assemblies, we conducted a comprehensive comparison against the start-of-the-art binary classifiers, including the viral classifiers DeepVirFinder and VIBRANT, the plasmid classifiers PlasClass and Platon, and the eukaryote classifiers EukRep, Whokaryote, and Tiara. Figure 1 summarizes the results. 4CAC outperforms almost all binary classifiers in each class classification, except in the classification of eukaryotes, where Tiara achieves a slightly higher F1 score on the Sim LF dataset (here and throughout, results were evaluated by their F1 score. See Methods for details). In classifying viruses, the XGBoost classifier designed in this study, without using the graph information, outperforms the start-of-the-art viral classifiers. In plasmid classification, the XGBoost classifier achieves the second-best performance in long-read assemblies, while Platon is the second-best in short-read assemblies. In classifying eukaryotes, all classifiers have good performance in long-read assemblies with Tiara and 4CAC achieving the best result. However, in short-read assemblies, 4CAC and the XGBoost classifier maintain consistently high F1 scores while the performance of the other eukaryote classifiers is markedly lower. Not surprisingly, by utilizing the graph information, 4CAC improved the XGBoost classification results in 11 out of 12 classifications across all datasets, and the improvement is dramatic in classifying plasmids from short-read assemblies.

**Fig. 1.**
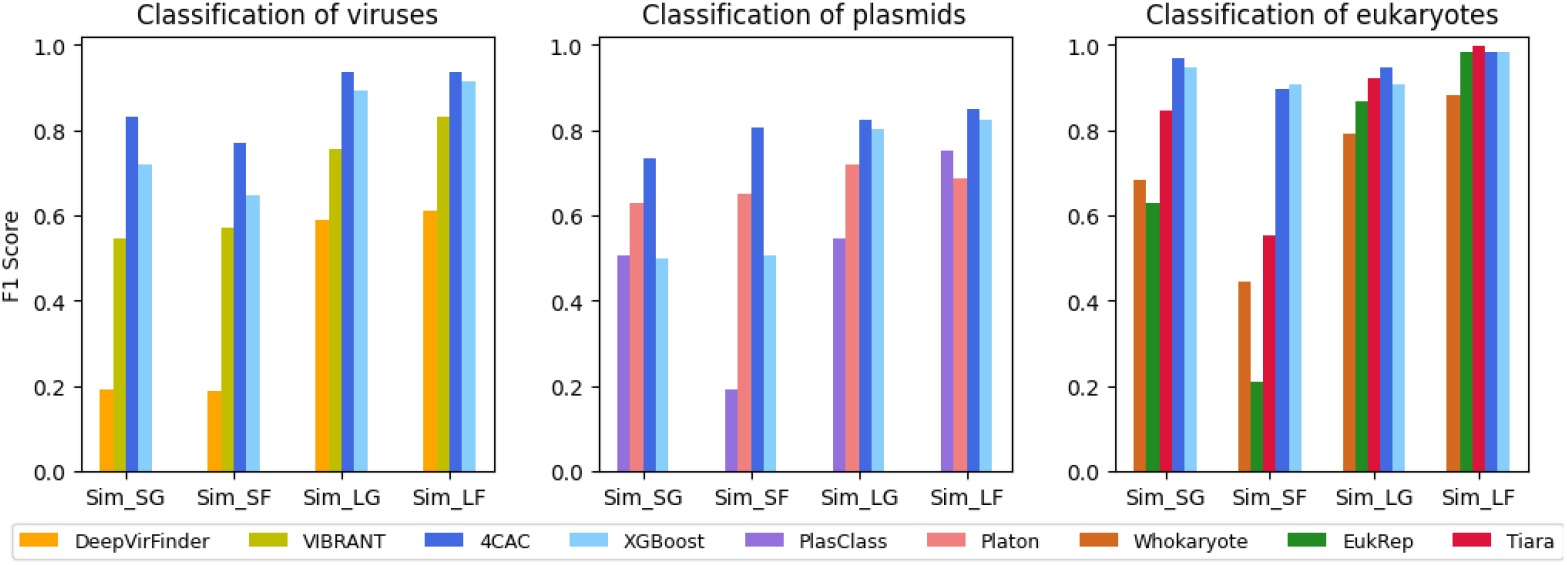
Performance of binary classifiers and 4CAC on simulated metagenomes. XGBoost represents the XGBoost classifier designed in this study without using graph information.

In addition, we conducted a comprehensive comparison between 4CAC and a set of threeway classifiers specifically designed to classify viruses and plasmids simultaneously from metagenome assemblies. The evaluated classifiers included PPR-Meta, viralVerify, geNomad, and 3CAC. Figure 2 summarizes the results. Across the various datasets, 4CAC consistently achieved the highest F1 scores in classifying both viruses and plasmids, outperforming the other classifiers. The only exception was observed in the Sim LF dataset, where 3CAC exhibited a slightly higher F1 score than 4CAC. 3CAC performed as the second-best classifier in most tests as it also utilizes graph information to improve its classification. Note that 3CAC utilizes either PPR-Meta or viralVerify to generate its classification. Thus, hereafter when using 3CAC, we executed the algorithm using both viralVerify and PPR-Meta solutions, and selected the better result. Among the stand-alone three-way classifiers, geNomad and PPR-meta had the highest F1 score in classifying viruses while viralVerify was the best in classifying plasmids in most tests.

**Fig. 2.**
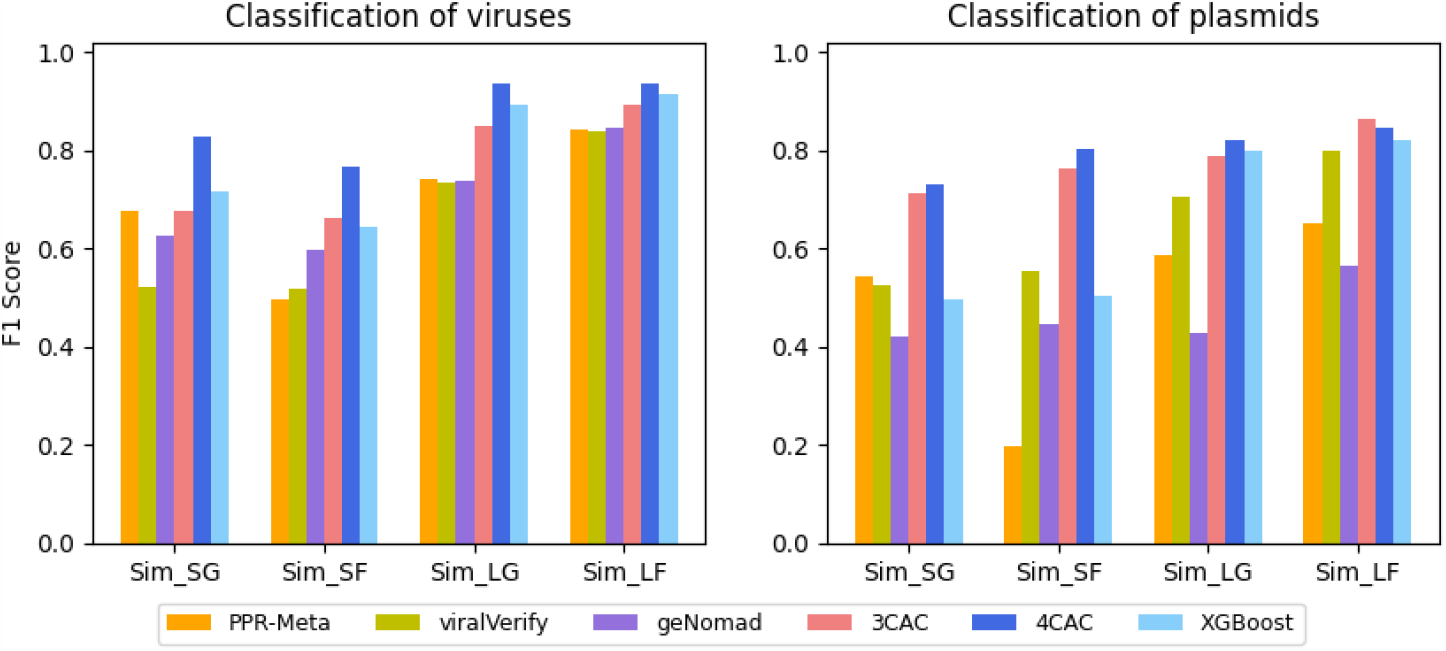
Performance of three-way classifiers and 4CAC on simulated metagenomes. XGBoost represents the XGBoost classifier designed in this study without using graph information.

It is important to note that, in order to ensure a fair comparison, eukaryotic contigs were excluded from our benchmark of three-way classifiers. Similarly, only two classes of contigs were considered when benchmarking binary classifiers. Supplementary Figures S1 and S2 provide a comprehensive overview of the results when all contigs were included. As expected, the inclusion of all contigs led to a decline in the performance of both binary and three-way classifiers, as they tend to misclassify contigs that are not modeled. For example, eukaryotic contigs and plasmid contigs can be misclassified as viruses by viral classifiers, and eukaryotic contigs can be misclassified as plasmids or viruses by three-way classifiers, etc. This highlights the need for a four-way classifier that is able to identify viruses, plasmids, eukaryotes, and prokaryotes simultaneously from metagenome assemblies.

### 2.4 Combining existing classifiers to generate four-way classifications

There are currently no four-class classifiers that can be compared with 4CAC. (A recent preprint reported a five-way classifier, DeepMicrobeFinder [13], but we could not run the code provided on GitHub.) Therefore, we combined existing classifiers to generate a fourway classification as follows. (1) The most straightforward idea is using VIBRANT and Platon to identify viruses and plasmids from metagenome assemblies. The remaining contigs are further classified as eukaryotes, prokaryotes, or uncertain by Tiara. We ran either VIBRANT or Platon first and selected the solution with a higher F1 score. This result is denoted by ***Binary+***. Here VIBRANT, Platon, and Tiara were selected because they performed best in binary classifications of viruses, plasmids, and eukaryotes from metagenome assemblies (shown in Figure 1). (2) Comparing three-way classifiers to binary classifiers demonstrated that 3CAC outperforms all binary classifiers in classifying viruses and plasmids from metagenome assemblies (Figure 1 and 2). Therefore, we further combined 3CAC with Tiara in the following way. We first classified contigs by 3CAC and set aside these classified as plasmids and viruses, then used Tiara to classify the rest into eukaryotes, prokaryotes, or uncertain. We also repeated the process in the reverse order, running first Tiara and then 3CAC. We then selected the solution with a higher F1 score. This result is denoted by ***3CAC+Tiara***.

Figure 3 demonstrates that 4CAC outperformed the combined classifiers in each classification across almost all datasets. In the long-read assembly Sim LF, 3CAC+Tiara had a slightly higher F1 score than 4CAC in classifying plasmids and eukaryotes. Compared to the initial XGBoost classification, 4CAC consistently improved the F1 score across all tests, and the improvement was more substantial in classifying plasmids from short-read assemblies. A possible reason is that plasmids often share similar sequences with their hosts, and contigs assembled from short reads are too short to accurately distinguish them. However, when considering the classification in the assembly graph, a chromosome contig that was misclassified as a plasmid is often surrounded by chromosome contigs. Thus, the graph refinement step of 4CAC can efficiently correct such a misclassification. The performance of combined classifiers exhibits greater variability across diverse datasets. Not surprisingly, 3CAC+Tiara outperformed Binary+ in almost all the tests. Compared to combined classifiers, 4CAC improved the F1 score remarkably in classifying eukaryotes and prokaryotes from short-read assembly Sim SF. This may be caused by a larger proportion of short contigs in Sim SF (58% in Sim SF vs. 19% in Sim SG. See Table 1). Short contigs are commonly unclassified by existing classifiers while 4CAC is able to classify most of them according to their neighboring long contigs in the assembly graph.

**Fig. 3.**
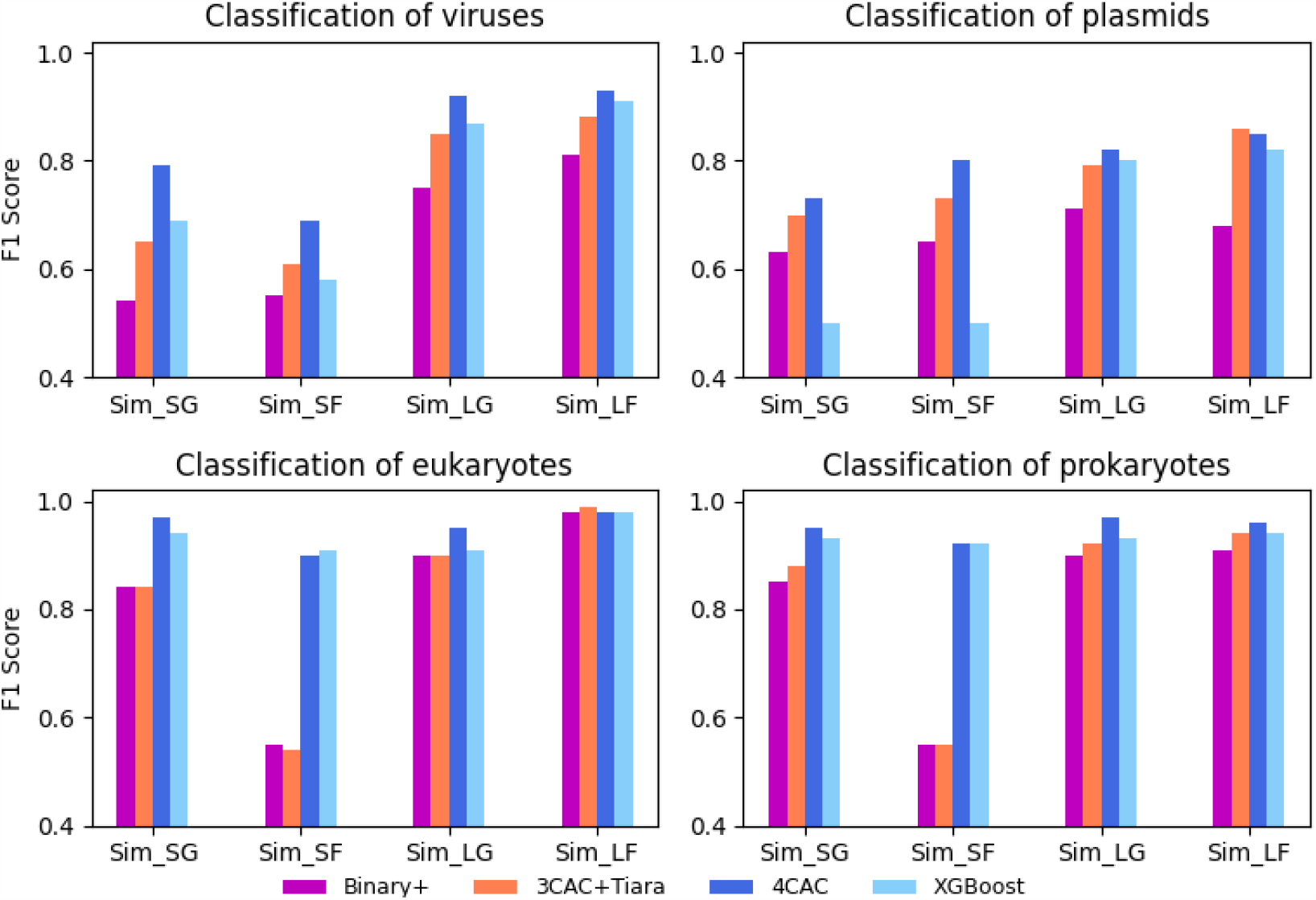
Performance of four-class classifiers on each class of simulated metagenomes. XGBoost represents the XGBoost classifier designed in this study without using graph information.

Figure 4 summarizes the total precision, recall, and F1 score of four-class classifiers. Consistent with the 3CAC algorithm, we observed that the graph refinement step improved the recall with little or no loss of precision in all the tests. 4CAC outperformed combined classifiers in both precision and recall in all the simulated assemblies, while XGBoost was the second-best. 4CAC improved the recall remarkably in Sim SF, due to a larger proportion of short contigs in it. Surprisingly, the XGBoost classifier itself, without using the graph information, had comparable or even better precision and recall than combined classifiers.

**Fig. 4.**
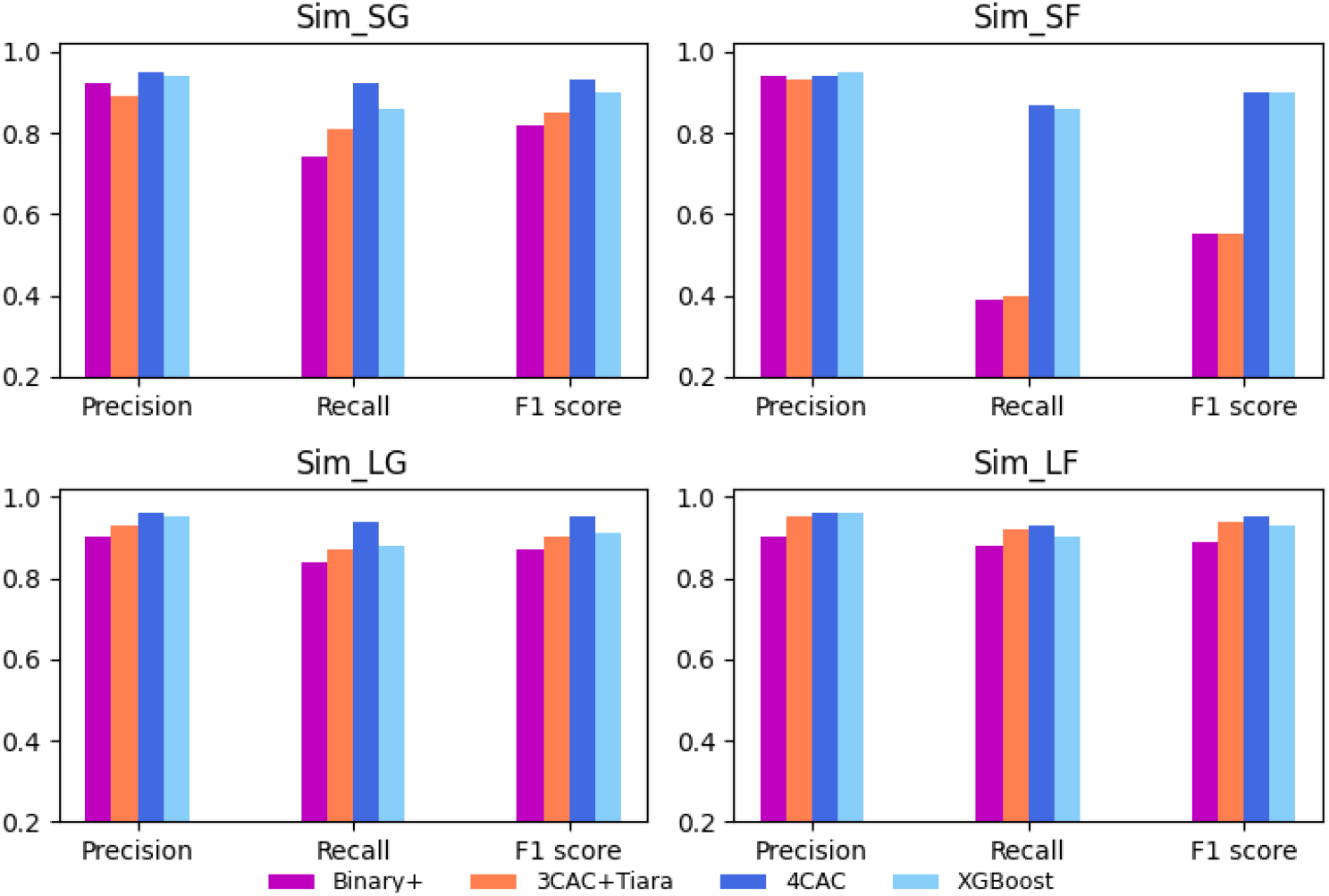
Performance of four-class classifiers on simulated metagenomes.

### 2.5 Performance on real metagenome samples

We additionally tested the performance of classifiers on four real complex metagenomic datasets: (1) Short-read sequencing of 18 preborn infant fecal microbiome samples (NCBI accession number SRA052203), referred to as **Sharon** [38]. (2) Short-read sequencing of a microbiome sample from the Tara Oceans (NCBI accession number ERR868402), referred to as **Tara** [15]. Currently, there is no study exploring microeukaryotes in long-read sequencing of microbiome samples. To test our method on long-read sequencing metagenomic datasets, we selected two publicly available datasets: (3) Oxford Nanopore sequencing of two human saliva microbiome samples (NCBI accession number DRR214963 and DRR214965), referred to as **Oral_Nano** [45]. (4) Pacbio HiFi sequencing of a human gut microbiome sample (NCBI accession number SRR15275211), referred to as **Gut_HiFi** [31]. Datasets with short reads and long reads were assembled by metaSPAdes and metaFlye, respectively. In Sharon and Oral_Nano, the multiple samples in each dataset were co-assembled. To identify the class of contigs in these real metagenome assemblies, we used all the complete assemblies of bacteria, archaea, viruses, plasmids, and microeukaryotes from the NCBI GenBank database as reference genomes and mapped contigs to these reference genomes using minimap2 [19]. A contig was considered matched to a reference sequence if it had ≥ 80% mapping identity along ≥ 80% of the contig length. Contigs that matched to reference genomes of two or more classes were excluded to avoid ambiguity. In all assemblies, contigs shorter than 500bp were not classified and excluded from the performance evaluation. Table 1 summarizes the properties of the datasets and the assemblies.

Since 3CAC+Tiara consistently outperformed the combination of binary classifiers (Figure 4), here we only compared 4CAC and its initial XGBoost classification to 3CAC+Tiara. Similar to the result in simulated assemblies, Figure 5 shows that the graph refinement step improved both the precision and recall of the XGBoost classification and led to significant improvement in the F1 score in most tests. In the Gut_HiFi dataset, 4CAC slightly improved the recall of XGBoost classification while sacrificing a few precision, and resulted in a similar F1 score. On the short read datasets Sharon and Tara, in which microeukaryotes were previously identified [41, 15], 4CAC achieved moderately better precision than 3CAC+Tiara but dramatically improved the recall. For example, 4CAC improved the recall from 0.54 to 0.87 in the Tara dataset. As a result, 4CAC had a substantially higher F1 score than 3CAC+Tiara.

**Fig. 5.**
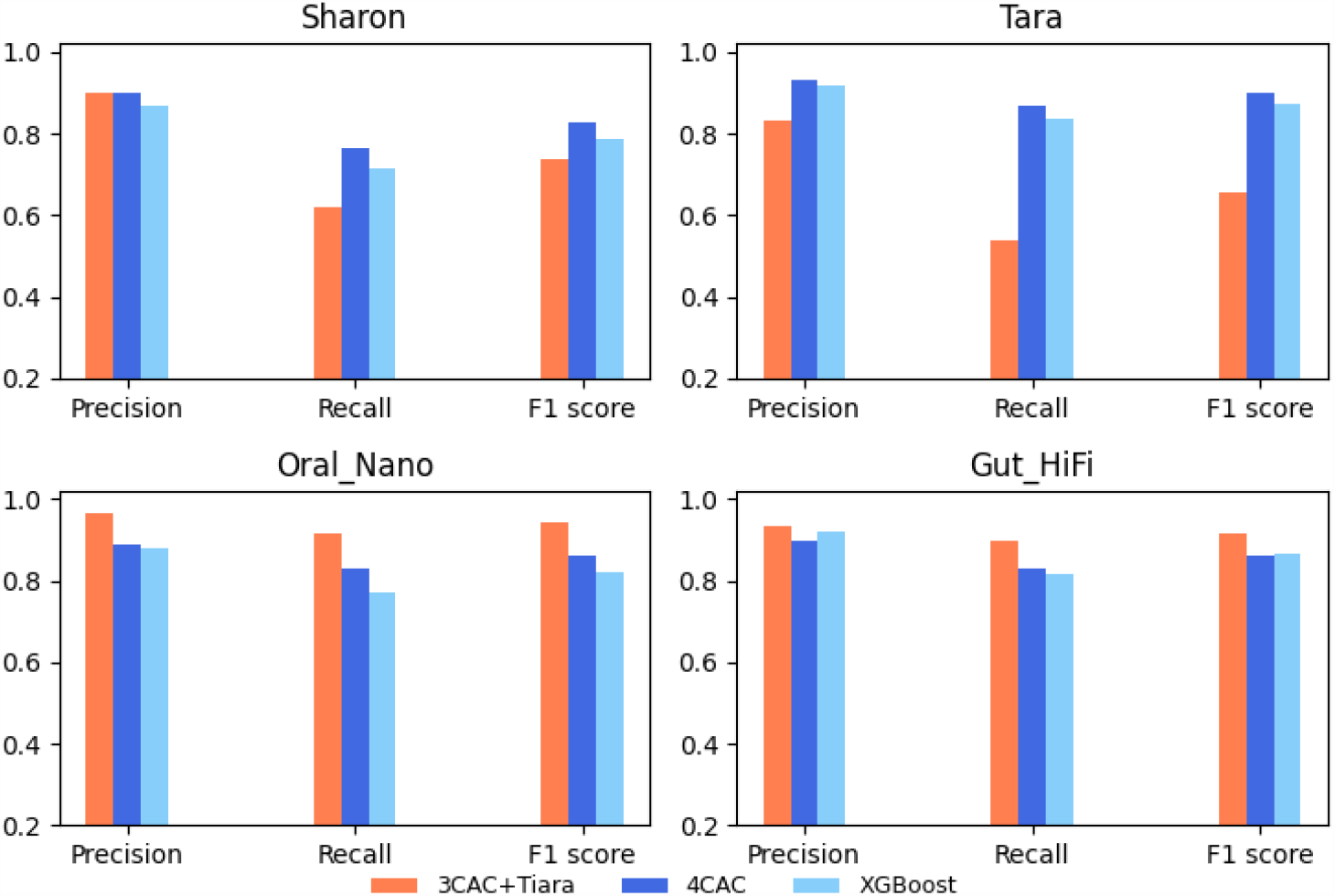
Performance of four-class classifiers on the real datasets. (a) Sharon and (b) Tara were assembled from short reads, (c) Oral_Nano and (d) Gut_HiFi were assembled from long reads.

Further analysis of the performance on the Sharon dataset reveals that the graph refinement step of 4CAC improved both the precision and recall of the XGBoost in each class classification (Figure 6). The improvement is more significant in the classification of plasmids, which is consistent with the observation on simulated assemblies [9]. Compared to 3CAC+Tiara, 4CAC had higher F1 scores in the classification of prokaryotes and eukaryotes, but a lower F1 score on viruses (Figure 6). A possible reason is that the proportion of viral contigs in the Sharon dataset is very small (0.6% vs. ≥ 1.3% in simulated assemblies. See Table 1). In this extreme case, viralVerify, which is used in 3CAC and classifies contigs based on their gene content, achieved higher precision than the machine learning-based methods, such as PPR-Meta and the XGBoost classifier.

**Fig. 6.**
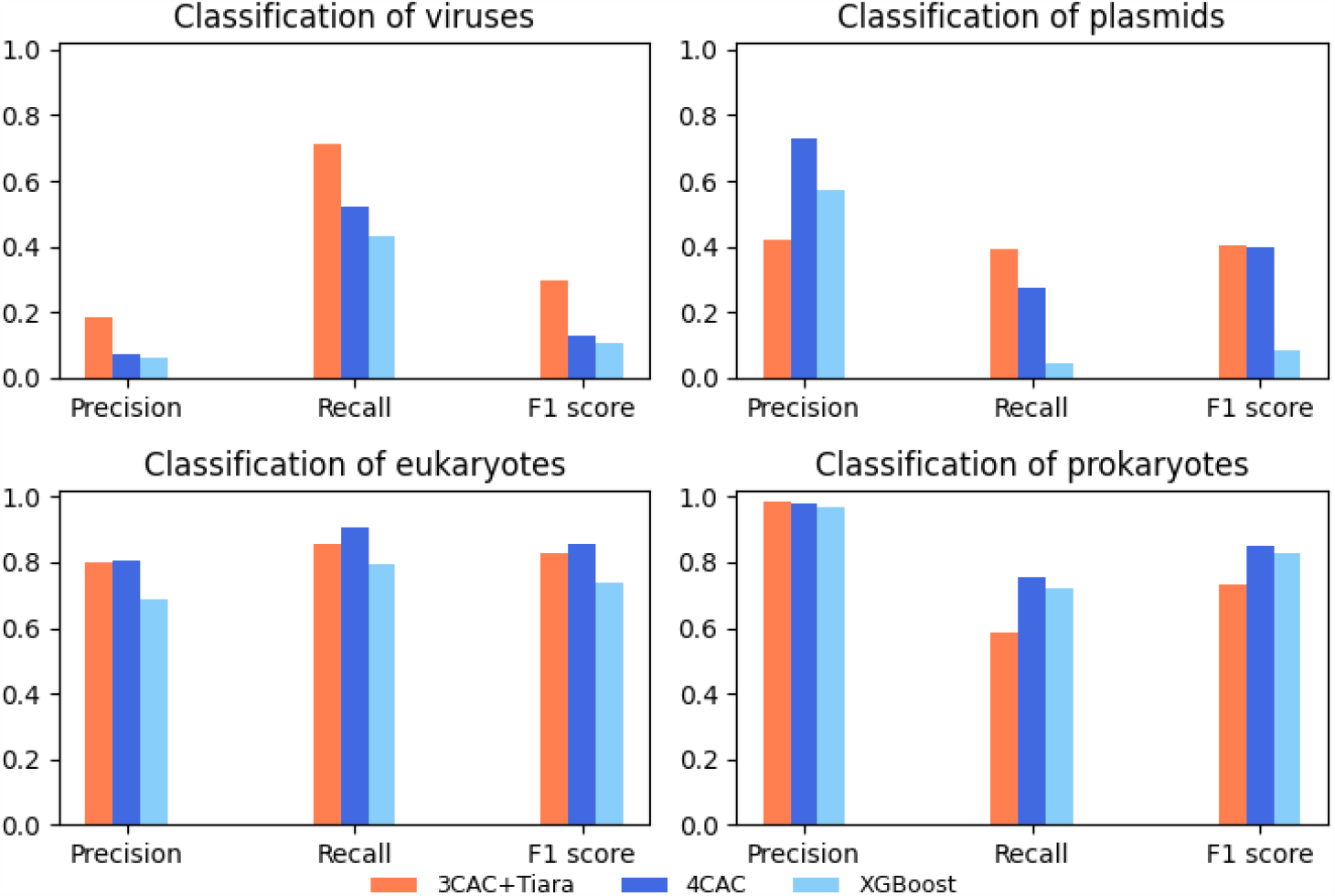
Performance on each class for real short read dataset Sharon.

On the two long-read datasets of human saliva and gut microbiome, 3CAC+Tiara outperformed 4CAC (Figures 5 (c) and (d)). Here too this is likely because each of the minor classes accounts for less than 0.6% of the contigs (Table 1).

## 3 Software and resource usage

Table 2 presents the runtime of the classifiers. All classifiers were run on contigs at least 500 bp in each dataset since contigs shorter than 500 bp were excluded from our evaluation. To run DeepVirFinder, we also excluded contigs longer than 2 Mb because DeepVirFinder failed on these long contigs. For 3CAC we report the runtime of viralVerify and PPR-Meta, since they required the lion’s share of the time, with the rest of 3CAC always taking less than 3 minutes. Due to the large runtime of viralVerify, geNomad, Platon, and VIBRANT, 4CAC is much faster than the other classifiers, which often require 1-2 orders of magnitudes more time. Supplementary Table S2 summarizes the memory usage of the classifiers. Memory usage was the highest for geNomad in all the tests. All runs were performed on a 44-core, 2.2GHz server with 792GB of RAM.

**Table 2.** Runtime of the tested classifiers. Runtime is measured by wall clock time in minutes. ViralV, PPR-M, and DeepVF represent classifiers viralVerify, PPR-Meta, and DeepVirFinder respectively.

|  | 4CAC | viralV | PPR-M | geNomad | Tiara | PlasClass | Platon | DeepVF | VIBRANT |
| --- | --- | --- | --- | --- | --- | --- | --- | --- | --- |
| Sim_SG | 6.9 | 322 | 60.4 | 106.9 | 2.9 | 4.2 | 241.5 | 91.2 | 185.2 |
| Sim_SF | 4.8 | 175 | 25.1 | 154.8 | 2.8 | 3.9 | 643.1 | 62.1 | 82.1 |
| Sim_LG | 3.7 | 185.4 | 33.4 | 92.8 | 1.3 | 2.3 | 22 | 33.2 | 139.8 |
| Sim_LF | 1.4 | 77.5 | 14.9 | 27.1 | 0.7 | 1.2 | 22 | 14.7 | 53.1 |
| Sharon | 1.4 | 29.9 | 7.3 | 16.4 | 0.5 | 1.2 | 49.4 | 16.3 | 25.3 |
| Tara | 14.3 | 221.1 | 94.7 | 155.8 | 4.4 | 11.2 | 638.9 | 92.3 | 50.6 |
| Oral_Nano | 8.2 | 452.5 | 84.4 | 140.1 | 3.5 | 5.6 | 301.1 | 88.6 | 201.1 |
| Gut_HiFi | 9.3 | 677.8 | 124 | 252.7 | 4.8 | 8.1 | 444.6 | 109.8 | 426.1 |

4CAC is freely available via https://github.com/Shamir-Lab/4CAC.

## 4 Discussion and Conclusion

We presented 4CAC, the first classification algorithm for simultaneously identifying viruses, plasmids, prokaryotes, and microeukaryotes in metagenome assemblies. Evaluation on simulated and real metagenomic datasets demonstrated that 4CAC substantially outperformed the combination of state-of-the-art binary and three-class classifiers in most tests. 4CAC also has a large speed advantage over the combined classifiers, running usually 1-2 orders of magnitude faster. In contrast to 3CAC, which necessitates the execution of viralVerify or PPR-meta, 4CAC is a stand-alone algorithm, making it more user-friendly.

Machine learning-based classifiers often assign scores to predictions, indicating their confidence. However, these scores do not reflect the true probabilities of the predictions. Indeed, when we attempted training XGBoost classifiers on class-imbalanced datasets using a default score threshold of 0.5 for all classes, results were unsatisfactory. By setting different probability thresholds for different classes, we obtained a good trade-off between precision and recall. Note, however, that when applying the same classifier to samples with varying class compositions, the results may exhibit significantly different false positive rates, and this is true for 4CAC as well.

On two real datasets assembled from long reads, where the relative abundance of viruses, plasmids, and eukaryotes was extremely low (less than 0.6% compared to over 1.3% in other assemblies), the combined classifier 3CAC+Tiara outperformed 4CAC. This difference in performance could potentially be attributed to the tendency of classifiers trained on *k*-mer compositions to yield a higher false positive rate compared to classifiers trained on the gene content of contigs. It is important to mention that these results may be biased by the underrepresentation of these classes in genomic databases. Given the current knowledge about species in metagenomes, we recommend using 4CAC on short reads and on host-filtered long read samples. For generic long read samples, where prokaryotes constitute the majority, we suggest utilizing 3CAC followed by Tiara for optimal results.

The implementation of the correction and propagation steps on the assembly graph yielded substantial improvements in the classification of short contigs. As anticipated, considering that 3CAC utilizes similar refinement procedures, the combined classifier 3CAC+Tiara demonstrated the second-best performance across all tests.

Our study has several limitations. First, as mentioned above, performance is affected by the relative abundance of the different classes in the input data. Second, the refinement step in 4CAC may misclassify some sequences, especially those that underwent horizontal gene transfer across classes, e.g. proviruses and integrated plasmids. However, as we have shown, that step improves overall performance. In future work, we aim to incorporate factors such as contig coverage and length to enhance the identification of proviruses.

Third, 4CAC does not categorize contigs at various taxonomic levels such as genus and species. Taxonomic classification requires different tools and approaches that are specifically designed for that goal, such as Kraken2 [42] and MetaPhlAn4 [5].

Finally, there is a chance of leakage occurring between the training and test set in case very similar sequences reside in both. However, using the yardstick of geNomad [8], where sequences with similarity *>* 95% are considered highly similar, we checked our test set and found that only *<* 15% of the test sequences share a high similarity with sequences in the training set. Moreover, the partition into training and test sets by GenBank release date is a common practice, which was also adopted in most of the classifiers that we evaluated (e.g. [9, 30, 35]). Furthermore, it also gives a realistic performance estimate, since when a method is applied to a new sample, some of the sequences encountered are likely to have highly similar counterparts in the database.

## 5 Methods

### 5.1 Training and testing datasets

To train and test the XGBoost classifier, we downloaded all complete assemblies of viruses, plasmids, prokaryotes (bacteria and archaea), and microeukaryotes (fungi and protozoa) from the National Center for Biotechnology Information (NCBI) GenBank database (download date April 22, 2022). After filtering out duplicate sequences, this database contained 31,129 prokaryotes, 69,882 viruses, 28,702 plasmids, and 2,486 microeukaryotes. To evaluate the ability of 4CAC to identify novel species, 24,734 prokaryotes, 65,475 viruses, 21,304 plasmids, and 2,315 microeukaryotes released before December 2021 were used to build the training set, while the remainder was used to build the testing set.

### 5.2 Training the XGBoost classifier

Inspired by previous studies [9, 30, 34], we trained several XGBoost models for different sequence lengths to assure the best performance. Specifically, five groups of fragments with lengths 0.5kb, 1kb, 5kb, 10kb, and 50kb were sampled from the reference genomes as artificial contigs. The composition information of each fragment is summarized by concatenating the canonical *k*-mer frequencies for *k* from 3 to 7, which results in a feature vector of length 10,952. We sampled 180k, 180k, 90k, 90k, and 50k fragments from each class to train the XGBoost models for sequence lengths 0.5kb, 1kb, 5kb, 10kb, and 50kb, respectively.

### 5.3 Length-specific classification

To assure the best classification for sequences of different lengths, we classify a sequence using the XGBoost model that is trained on fragments with the most similar length. Namely, the five XGBoost models we trained above are used to classify sequences in the respective length ranges (0, 0.75kb], (0.75kb, 3kb], (3kb, 7.5kb], (7.5kb, 30kb], and (30kb, ∞]. Given a sequence, we calculate its canonical *k*-mer frequency vector and use it as the feature vector to classify the sequence with the model that matches its length. The calculation of *k*-mer frequency vector can be performed in parallel for different sequences to achieve faster runtime.

For each sequence in the input, the XGBoost classifier outputs four scores between 0 and 1 indicating its confidence of being classified as a virus, plasmid, prokaryote, or microeukaryote. Taking into consideration the class imbalance, we chose to set different thresholds for classifying different classes. Tests on simulated metagenomes show that increasing score thresholds for prokaryotes and eukaryotes had little effect on the precision but decreased the recall a lot (Supplementary Figure S3). Thus we did not set specific score thresholds for prokaryotes and eukaryotes. In other words, a sequence was classified as prokaryote or eukaryote if that class had the highest score, irrespective of its value. For viruses and plasmids, we tested several score thresholds (0.8, 0.85, 0.9, 0.95) and similar results were observed, while increasing the score threshold slightly improved the result in both precision and recall (see Supplementary Table S1). Note that increasing the score threshold did not decrease the recall of 4CAC, because the graph refinement step can significantly improve the recall over the initial classification. Therefore, in the 4CAC algorithm, we set the default score threshold of 0.95 for classifying contigs as viruses and plasmids.

### 5.4 Refining the classification using the assembly graph

Nodes in an assembly graph represent contigs and edges represent sequence overlaps between the corresponding contigs. In our description below, the neighbors of a contig are its adjacent nodes in the undirected assembly graph. In the 4CAC algorithm, we exploit the assembly graph to improve the initial classification by the following two steps. The description here follows [33].

1. Correction of classified contigs. All classified contigs are scanned in decreasing order of the number of their classified neighbors. For a classified contig *c*, if it has at least two classified neighbors and all of them belong to the same class while *c* belongs to a different class, 4CAC corrects the classification of *c* to be the same as its classified neighbors. Note that once a contig was corrected, the class of this contig and its classified neighbors will not be corrected anymore.
2. Propagation to unclassified contigs. For an unclassified contig *c*, if all of its classified neighbors belong to the same class, 4CAC assigns *c* to that class. Unclassified contigs are scanned and classified in decreasing order of the number of their classified neighbors. We repeat this step until no propagation is possible.

### 5.5 Simulated datasets

We randomly selected 100 prokaryotes, 461 co-existing plasmids, 500 viruses, and 6 microeukaryotes from the NCBI GenBank Database to mimic species in a microbial community. All the genomes selected were released after December 2021, and thus they were not included in the training set of the classifier. Two short-read and two long-read metagenome assemblies were generated from this microbial community as follows. As a *generic metagenome* scenario, we simulated reads from prokaryotes, eukaryotes, viruses, and plasmids in a ratio of 56:24:10:10. As a *filtered metagenome* scenario, where reads from large genomes are filtered and thus plasmids and viruses are enriched, we simulated reads from prokaryotes, eukaryotes, viruses, and plasmids in a ratio of 14:6:40:40. The abundance profiles of prokaryotes, eukaryotes, and viruses were modeled by the log-normal distribution as in [30]. The copy numbers of co-existing plasmids were simulated by the geometric distribution with parameter *p* = *min*(1, *log*_10_(*L*)*/*7) where *L* is the plasmid length as in [30]. The abundance profile of plasmid genomes was calculated from their host abundance profile and the copy numbers of plasmids. 150bp short reads were simulated from the genome sequences using InSilicoSeq and assembled by metaSPAdes. Long reads were simulated from the genome sequences using NanoSim and assembled by metaFlye. The error rate of long reads was 9.8% and their average length was 14.9kb. For each assembly, contigs were mapped to the reference genomes by metaQUAST to define the ground truth. To ensure confident assignment of contigs, metaQUAST excludes contigs shorter than 500bp by default.

### 5.6 Evaluation criteria

All the classifiers were evaluated based on precision, recall, and F1 scores calculated as follows.

– **Precision:** the fraction of correctly classified contigs among all classified contigs. Note that uncertain contigs were not included in this calculation.
– **Recall:** the fraction of correctly classified contigs among all contigs.
– **F1 score:** the harmonic mean of the precision and recall, or equivalently: *F* 1 *score* = (2 ∗ *precision* ∗ *recall*)*/*(*precision* + *recall*).

Following [30, 9], the precision, recall, and F1 scores here were calculated by counting the number of contigs and did not take into account their length. The precision and recall were also calculated separately for virus, plasmid, prokaryote, and eukaryote classification. For example, the precision of virus classification was the fraction of correctly classified virus contigs among all contigs classified as viruses, and the recall of virus classification was the fraction of correctly classified virus contigs among all virus contigs.

## Declarations

## Acknowledgments

We thank members of the Shamir Lab for their help and comments - David Pellow, Ron Saad, and Hagai Levi.

## Availability of data and materials

4CAC is implemented in Python and is available on GitHub (https://github.com/Shamir-Lab/4CAC). All sequencing datasets analyzed in this study are available in National Center for Biotechnology Information (NCBI), accession numbers: SRA052203 for the Sharon dataset, ERR868402 for the Tara Ocean dataset, DRR214963 and DRR214965 for the Oral_Nano dataset, and SRR15275211 for the Gut_HiFi dataset.

## Ethics approval and consent to participate

Not applicable.

## Consent for publication

Both authors consented.

## Authors’ contributions

LP and RS designed the study. LP developed and implemented the 4CAC algorithm, analyzed the datasets, and wrote the manuscript. RS oversaw the development of the project and edited the manuscript. Both authors read and approved the final manuscript.

