## Supplementary files for "4CAC: 4-class classifier of metagenome contigs using machine learning and assembly graphs"

### 4CAC: 4-class classification of metagenomic assemblies using machine learning and assembly graphs

#### Supplementary Material

#### 1 Supplementary tables

**Table S1. Performance of 4CAC on simulated metagenomes with initial classifications generated by different score thresholds.** To generate an initial four-way classification by XGBoost classifier, we tested score thresholds 0.8, 0.85, 0.9, and 0.95 to classify contigs as phages and plasmids. Note that this table presents the results of the full 4CAC algorithm.

| Datasets | Score | Phage | | | Plasmid | | | Prokaryote | | | Eukaryote | | | All | | |
|---|---|---|---|---|---|---|---|---|---|---|---|---|---|---|---|---|
| | threshold | precision | recall | F1 score | precision | recall | F1 score | precision | recall | F1 score | precision | recall | F1 score | precision | recall | F1 score |
| | 0.95 | 0.79 | 0.79 | 0.79 | 0.74 | 0.66 | 0.70 | 0.96 | 0.94 | 0.95 | 1.00 | 0.95 | 0.97 | 0.95 | 0.92 | 0.93 |
| Sim_SG | 0.9 | 0.70 | 0.84 | 0.77 | 0.65 | 0.75 | 0.69 | 0.96 | 0.92 | 0.94 | 1.00 | 0.94 | 0.97 | 0.93 | 0.91 | 0.92 |
| | 0.85 | 0.64 | 0.87 | 0.73 | 0.60 | 0.79 | 0.68 | 0.97 | 0.91 | 0.94 | 1.00 | 0.93 | 0.96 | 0.92 | 0.91 | 0.91 |
| | 0.8 | 0.60 | 0.88 | 0.71 | 0.56 | 0.83 | 0.67 | 0.97 | 0.89 | 0.93 | 1.00 | 0.93 | 0.96 | 0.91 | 0.90 | 0.91 |
| | 0.95 | 0.60 | 0.80 | 0.69 | 0.82 | 0.76 | 0.79 | 0.94 | 0.90 | 0.92 | 0.98 | 0.85 | 0.91 | 0.95 | 0.88 | 0.91 |
| Sim_SF | 0.9 | 0.42 | 0.85 | 0.56 | 0.63 | 0.81 | 0.71 | 0.94 | 0.90 | 0.92 | 0.98 | 0.85 | 0.91 | 0.94 | 0.88 | 0.90 |
| | 0.85 | 0.32 | 0.87 | 0.47 | 0.50 | 0.84 | 0.63 | 0.94 | 0.89 | 0.91 | 0.98 | 0.85 | 0.91 | 0.92 | 0.87 | 0.90 |
| | 0.8 | 0.25 | 0.89 | 0.39 | 0.40 | 0.87 | 0.55 | 0.94 | 0.87 | 0.91 | 0.98 | 0.85 | 0.91 | 0.90 | 0.86 | 0.88 |
| | 0.95 | 0.91 | 0.91 | 0.91 | 0.84 | 0.79 | 0.82 | 0.98 | 0.97 | 0.97 | 1.00 | 0.90 | 0.95 | 0.96 | 0.94 | 0.95 |
| Sim_LG | 0.9 | 0.90 | 0.92 | 0.91 | 0.79 | 0.83 | 0.81 | 0.98 | 0.96 | 0.97 | 1.00 | 0.90 | 0.95 | 0.95 | 0.94 | 0.95 |
| | 0.85 | 0.86 | 0.93 | 0.89 | 0.76 | 0.89 | 0.82 | 0.98 | 0.95 | 0.96 | 1.00 | 0.90 | 0.95 | 0.95 | 0.94 | 0.94 |
| | 0.8 | 0.84 | 0.93 | 0.88 | 0.70 | 0.92 | 0.79 | 0.99 | 0.93 | 0.96 | 1.00 | 0.90 | 0.95 | 0.94 | 0.93 | 0.93 |
| | 0.95 | 0.96 | 0.90 | 0.93 | 0.88 | 0.81 | 0.84 | 0.97 | 0.95 | 0.96 | 1.00 | 0.97 | 0.98 | 0.96 | 0.93 | 0.95 |
| Sim_LF | 0.9 | 0.93 | 0.91 | 0.92 | 0.85 | 0.85 | 0.85 | 0.97 | 0.95 | 0.96 | 1.00 | 0.97 | 0.98 | 0.95 | 0.93 | 0.94 |
| | 0.85 | 0.91 | 0.91 | 0.91 | 0.78 | 0.90 | 0.84 | 0.97 | 0.93 | 0.95 | 1.00 | 0.97 | 0.98 | 0.94 | 0.93 | 0.94 |
| | 0.8 | 0.89 | 0.92 | 0.90 | 0.76 | 0.92 | 0.83 | 0.98 | 0.92 | 0.95 | 1.00 | 0.97 | 0.98 | 0.94 | 0.93 | 0.93 |

**Table S2. Memory usage of the tested classifiers in GB.** ViralV, PPR-M, and DeepVF represent classifiers viralVerify, PPR-Meta, and DeepVirFinder respectively.

| | **4CAC** | **viralV** | **PPR-M** | **geNomad** | **Tiara** | **PlasClass** | **Platon** | **DeepVF** | **VIBRANT** |
|---|---|---|---|---|---|---|---|---|---|
| Sim_SG | 3.9 | 1.3 | 6.5 | 78.1 | 2.2 | 5.7 | 14.7 | 21.3 | 0.2 |
| Sim_SF | 15.5 | 0.7 | 6.4 | 66.4 | 1.6 | 16.9 | 14.8 | 10.2 | 0.2 |
| Sim_LG | 0.7 | 0.2 | 6.3 | 66.7 | 1.4 | 0.5 | 12.2 | 13.8 | 0.2 |
| Sim_LF | 0.6 | 0.3 | 6.3 | 68.2 | 1.4 | 0.4 | 12.1 | 7.9 | 0.2 |
| Sharon | 9.9 | 0.1 | 6.3 | 65.7 | 1.4 | 7.3 | 14 | 9.3 | 0.1 |
| Tara | 26.2 | 9.8 | 12.3 | 67.9 | 2.8 | 25.2 | 15.9 | 2.8 | 0.1 |
| Oral_Nano | 2.2 | 1.5 | 6.2 | 55.4 | 2.8 | 2.9 | 15 | 10 | 0.2 |
| Gut_HiFi | 2.7 | 2.4 | 9.3 | 68.9 | 3.7 | 3.3 | 13.2 | 9.4 | 0.3 |

#### 2 Supplementary figures

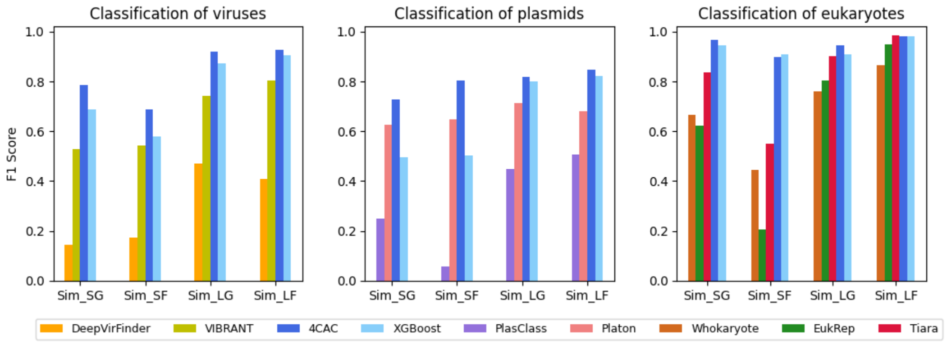


**Fig. S1. Performance of binary classifiers and 4CAC on simulated metagenomes** All four classes of contigs were included in the benchmark.

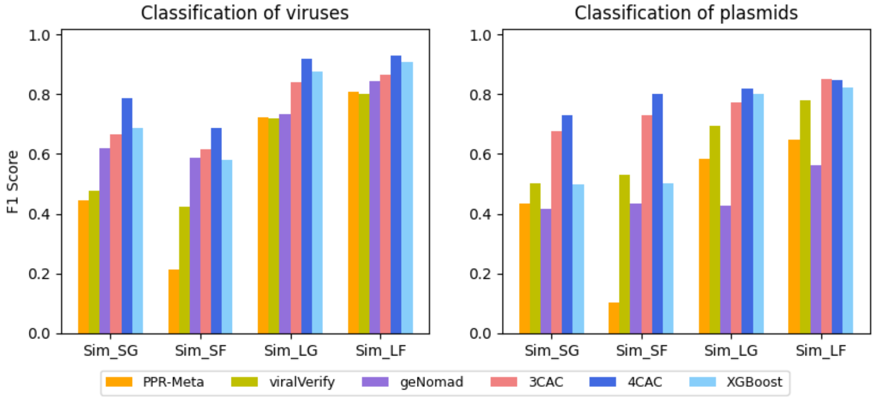


**Fig. S2. Performance of three-way classifiers and 4CAC on simulated metagenomes** All four classes of contigs were included in the benchmark.

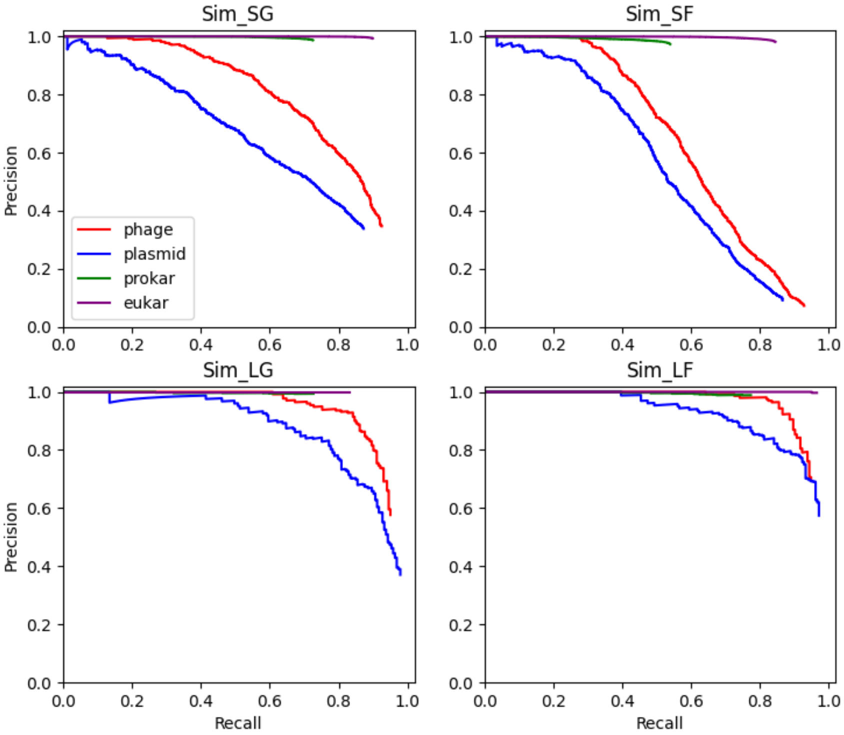


**Fig. S3. The precision-recall curve of our XGBoost classifier on simulated metagenomes.** Lines colored red, blue, green, and purple represent precision-recall curves for phage, plasmid, prokaryote, and eukaryote classification, respectively. Sim_SG and Sim_SF are datasets of short reads while Sim_LG and Sim_LF are datasets of long reads.
